# An ultrasensitive, rapid, and portable coronavirus SARS-CoV-2 sequence detection method based on CRISPR-Cas12

**DOI:** 10.1101/2020.02.29.971127

**Authors:** Curti Lucia, Pereyra-Bonnet Federico, Gimenez Carla Alejandra

**Affiliations:** INPA-National Scientific and Technical Research Council (CONICET)- Argentina-University of Buenos Aires, Argentina; CASPR Biotech, San Francisco California, USA

**Keywords:** CRISPR-Cas12, emerging virus, SARS-CoV2, COVID-19, 2019-nCoV, diagnosis

## Abstract

Severe acute respiratory syndrome coronavirus 2 (SARS-CoV-2) has received global attention due to the recent outbreak in China. In this work, we report a CRISPR-Cas12 based diagnostic tool to detect synthetic SARS-CoV-2 RNA sequences in a proof-of-principle evaluation. The test proved to be sensitive, rapid, and potentially portable. These key traits of the CRISPR method are critical for virus detection in regions that lack resources to use the currently available methods.

## Introduction

The rapid spread of the novel coronavirus is clearly a major concern for countries across the world. This virus is a single-stranded RNA virus with high sequence identity to SARS-CoV and has, therefore, been named SARS-CoV-2. This coronavirus variant is capable of transmission from person to person, which makes it a dangerous virus that can cause a pandemic.

Several SARS-CoV-2 detection assays have been reported to be currently under development. The WHO website provides information on several virus detection protocols that have been used in different countries such as China, Germany, Japan, and the US, among others (*1*). All these are real-time reverse transcription PCR (rRT-PCR) based assays, and despite their established efficiency, require highly specialized personnel and expensive equipment for implementation. In such a backdrop, any development toward ultrasensitive, cheaper, and portable diagnostic tests for the assessment of suspected cases, regardless of the presence of qualified personnel or sophisticated equipment for virus detection, could help advance the diagnosis of COVID-19.

CRISPR is a biotechnological technique well-known for its use in gene editing. Notably, CRISPR has been recently used for the *in vitro* detection of nucleic acids, thereby emerging as a powerful and precise tool for molecular diagnosis (*2-4*). Within the CRISPR-Cas effector family, Cas12 is a RNA-guided DNase belonging to the class II type V-A system that induces indiscriminate single-stranded DNA (ssDNA) collateral cleavage after target recognition. This leads to the degradation of ssDNA reporters that, emit a fluorescence signal on cleavage or alternatively, could be detected on a paper strip (by lateral flow) in a portable manner (5). Therefore, CRISPR-Cas12 based tools possess the potential to emerge as an *in situ* diagnostic tool for rapid detection of the SARS-CoV-2 virus. In this work, we employed CRISPR-Cas12a and its unspecific collateral ssDNAse activity to generate a fast, accurate, and portable SARS-CoV-2 sequence detection method.

## Materials and methods

For the detection assays, we included synthetic RNA fragments of SARS-CoV-2 corresponding to the *RdRp, ORF1b* and *ORF1ab* genes, using the WH-human1 sequence (GenBank MN908947) as a reference. Briefly, the SARS-CoV-2 fragments were synthesized as complementary DNA oligonucleotides and treated with fill-in PCR (NEBNext® High-Fidelity 2X PCR Master Mix) to generate the DNA templates. These DNA templates were transcribed into RNA using an *in vitro* transcription (IVT) Kit (Ambion, Invitrogen) under the control of a T7 promoter.

A target amplification step was performed using the TwistAmp® Basic recombinase polymerase amplification (RPA) kit (TwistDx, Cambridge, United Kingdom) and RT was carried out by the addition of 5 μL of synthetic SARS-CoV-2-RNA input, 2.5 μL of M-MuLV reverse transcriptase (NEB), and 1 µL of murine RNase inhibitor (NEB) in a 50 μL final reaction volume. Reactions were run for 30 min at 42°C. The RT-RPA was carried out in one step, with the same pair of primers for both reactions.

To generate the CRISPR-detection complex, we mixed 75 nM of the commercially available LbCas12a endonuclease (NEB) with the same amount of single guide RNA (sgRNA) synthesized in-house by hybridization of DNA oligonucleotides followed by fill-in PCR and IVT. The complexation reaction was carried out in a solution containing 1X NEBuffer 2.1 (NEB) (composed of 100 mM NaCl, 50 mM Tris-HCl, 10 mM MgCl_2_, 1 mM DTT, pH 7.9) at room temperature for 10 min.

For plate reader-based assays, ssDNA reporters labeled with FAM were included in the detection mix; while for portable detection with paper strips, ssDNA reporters labeled with biotin were included. The reaction was initiated by diluting the LbCas12a complexes to a final concentration of 37 nM LbCas12a : 37 nM sgRNA in a solution containing 1X NEBuffer 2.1 (NEB) and 1 µM custom ssDNA reporter substrates in a 40 μL reaction volume. A 2 μL aliquot of RT-RPA input was used. Reactions were incubated for up to 90 min at 37°C. Fluorescence measurements were acquired at 10 min intervals (λ_ex_: 485 nm and λ_em_: 535 nm) in a SpectraMax M2 fluorescence plate reader (Molecular Devices) operated in the 384-well microplate format. For paper-based measurements, we used the Milenia HybriDetect 1 (TwistDx) lateral flow system, as per the manufacturer’s instructions. Finally, we simulated a clinical sample by adding synthetic SARS-CoV-2 RNA fragments (10^5^ copies/µL) to saliva samples from a healthy donor and measuring the results on the test strips. The saliva was treated with heat and chemicals before using 2 µL as input for RT-RPA (5).

Details about the target sequences, sgRNA, primers for RT-RPA, and ssDNA reporters are listed in the accompanying table 1. The general scheme of target synthesis, amplification, detection with CRISPR, and visualization of the result are shown in the figure 1.

**Table.**
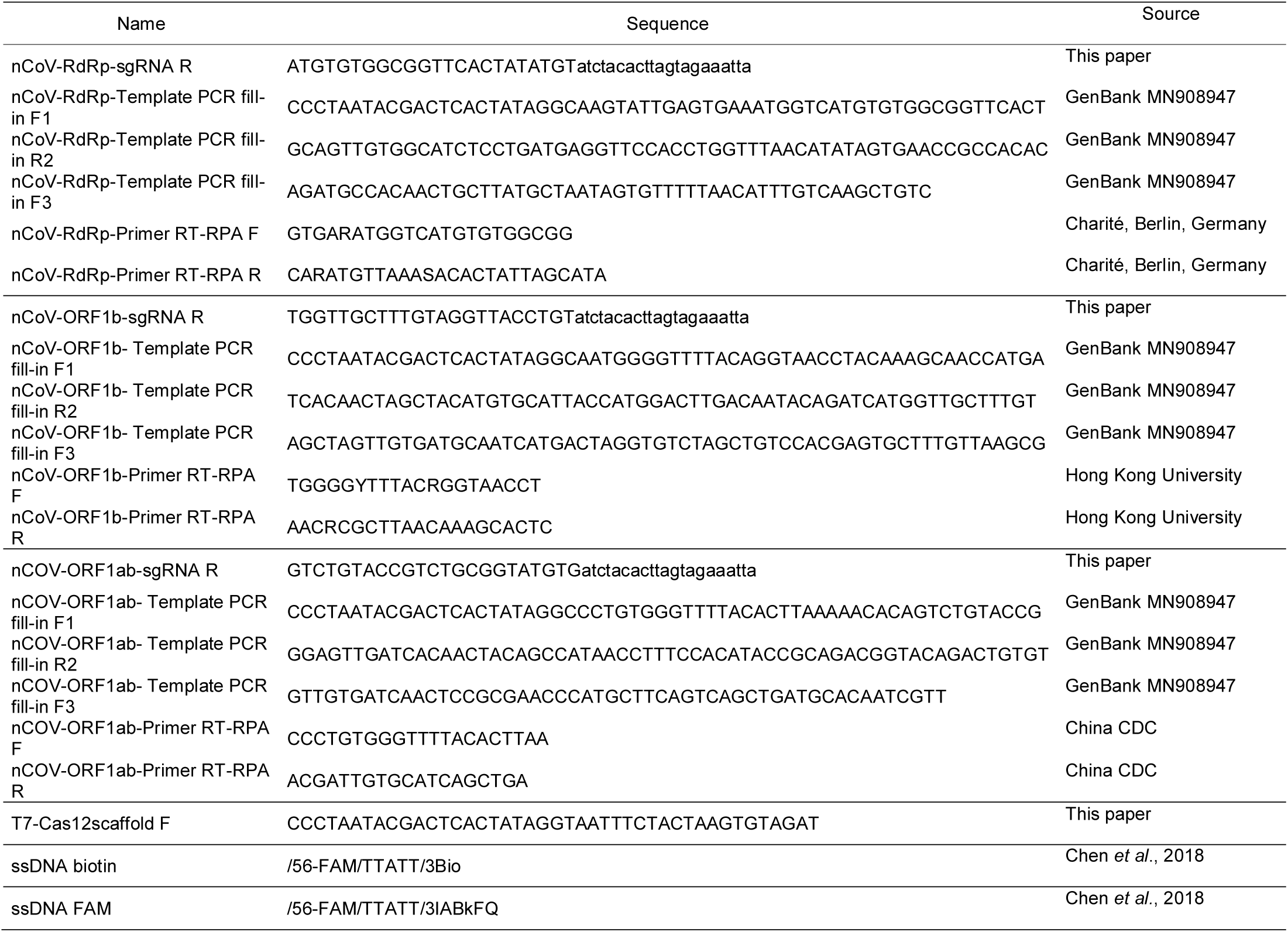
List of DNA oligonucleotides and ssDNA reporters used in this work.

**Figure 1.**
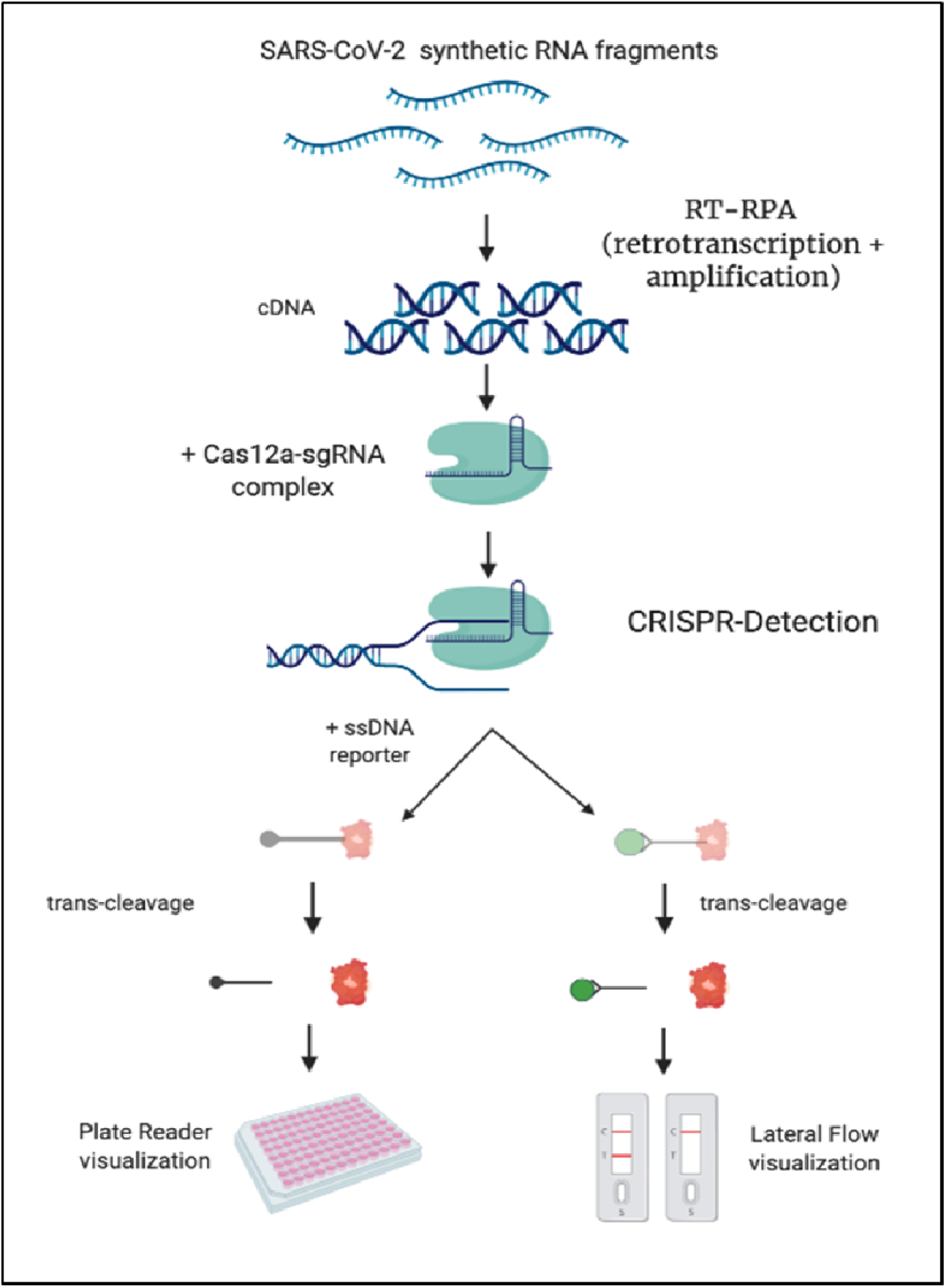
CRISPR-based detection method for novel coronavirus. General scheme of CRISPR detection procedure. In the two strategies all the process takes less than 60 min.

## Results and Discussion

The best results were obtained in attempts to detect the *ORF1ab* region with primers suggested by the China CDC (*6*). Based on visualization of these results in a fluorescence spectrophotometer, the limit of detection (LOD) for *ORF1ab* coronavirus sequences was estimated to be up to 10 copies/µL (figure 2 panel A of the accompanying figure), which is 4 orders of magnitude lower than the viral load found for the patient reported in Berlin (10^5^ copies/µL) (*7*).

**Figure 2.**
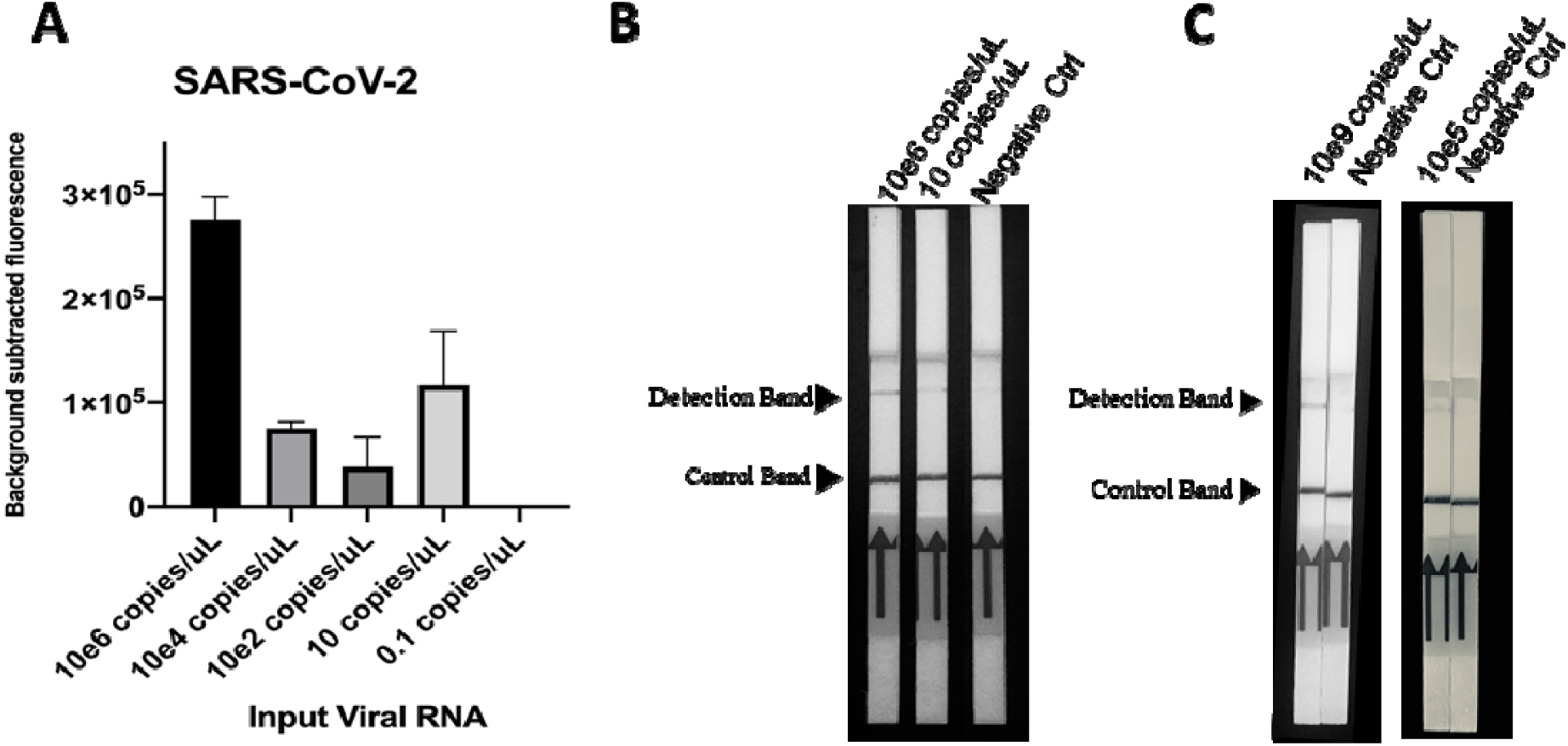
A) Assessment of the detection limit of synthetic SARS-CoV-2 by the CRISPR-based assay, inferred from fluorescence values after 30 min. Background subtracted fluorescence represents sample minus control fluorescence without target. Detection of synthetic SARS-CoV-2 RNA using commercially available paper strips (lateral-flow) in buffer (B) or in saliva (C). The optimization in saliva} continues in process to obtain more intense signal on paper strip. Microsoft Excel 2016 was used to arrange the data for analysis. GraphPad Prism v.8.1.2 (San Diego, USA) was used to plot the graph. In all cases, data represent mean ± SD (n=3).

When assays were performed using a paper strip, we obtained a similar LOD, demonstrating that the detection system could be rendered portable without loss in sensitivity (figure 2 panel B of the accompanying figure).

Finally, we simulated a clinical sample by adding synthetic SARS-CoV-2 RNA fragments to saliva samples collected from a healthy person. There are advantages using saliva for the diagnosis of COVID-19. Saliva specimens can be provided by the patient easily without any invasive procedures (8). Positive results for this artificial saliva-sample were obtained using the paper strip-based assay (figure 2 panel C). These latest results indicate that the CRISPR detection system reported here was not inhibited by naturally occurring molecules in the saliva and can reach real saliva specimens concentration (8), therefore, emerging to be a promising method got the rapid and portable detection of clinical cases of COVID-19.

## Conclusions

The current rRT-PCR-based COVID-19 diagnostic approaches have been shown to be very efficient and accurate for virus detection. However, the virus can spread to regions where the equipment required to perform rRT-PCR is not available. We demonstrated that the CRISPR-Cas12 based detection method is characterized by a LOD value lower than the minimum levels needed to presently detect the virus in clinical samples. The main advantage of the CRISPR-diagnostic method reported here is its portability and low cost (USD 1-2/reaction) (*2*). A possible criticism for this study is that the method has not been tested using patient samples. This is because clinical cases have not yet been reported in our region (South America). Nevertheless, our work represents a proof-of-principle study of the usefulness of the CRISPR-Cas12 detection technology that could be deployed worldwide without the requirement of sophisticated instrumentation.

## Note

While we were preparing this paper, another two protocols for SARS-CoV-2 detection using CRISPR diagnostics (SHERLOCK, v.20200214; DETECTER v.20200215) was published, confirming the potential of the technique.

## Acknowledgments

The authors thanks Guillermo Repizo from IBR-CONICET for his critical suggestion on this manuscript.

